# Canonical time-series features for characterizing biologically informative dynamical patterns in fMRI

**DOI:** 10.1101/2024.07.14.603477

**Authors:** Imran Alam, Brendan Harris, Patrick Cahill, Oliver Cliff, Marija Markicevic, Valerio Zerbi, Ben D. Fulcher

## Abstract

The interdisciplinary time-series analysis literature encompasses thousands of statistical features for quantifying interpretable properties of dynamical data. But for any given application, it is likely that just a small subset of informative time-series features is required to capture the dynamical quantities of interest. So, while comprehensive libraries of time-series features have been developed, it is useful to construct reduced and computationally efficient subsets for specific applications. In this work, we demonstrate a systematic process to deduce such a reduced set, focused on the problem of distinguishing changes to functional Magnetic Resonance Imaging (fMRI) time series caused by a range of experimental manipulations of excitatory and inhibitory neural activity in mouse cortical circuits. We reduce a comprehensive library of over 7000 candidate time-series features down to a subset of 16 features, which we call *catchaMouse16*, that aims to both: (i) accurately characterize biologically relevant properties of fMRI time series; and (ii) minimize inter-feature redundancy. The *catchaMouse16* feature set accurately classifies experimental perturbations of neuronal activity from fMRI recordings, and also shows strong generalization performance on an unseen mouse and human resting-state fMRI data where it tracks spatial variations in excitatory and inhibitory cortical cell densities, often with greater statistical power than the full *hctsa* feature set. We provide an efficient, open-source implementation of the *catchaMouse16* feature set in C (achieving an approximately 60 times speed-up relative to the native Matlab code of the same features), with wrappers for Python and Matlab. This work demonstrates a procedure to reduce a large candidate time-series feature set down to the key statistical properties of mouse fMRI dynamics that can be used to efficiently quantify and interpret informative dynamical patterns in neural time series.

## Introduction

The study of how systems evolve over time is ubiquitous across the sciences, with contributions to time-series analysis from fields ranging from biomedicine to astrophysics. Recent work has assembled these myriad techniques into a unified library of over 7000 time-series ‘features’—the *hctsa* feature set [1, 2]—where each feature encapsulates an interpretable (real-valued) property of a time series. Time-series features quantitatively capture different types of properties from the dynamics of a process, such as how periodic it is, how it distributes its energy across different frequency components, how predictable it is, how nonlinear the dynamics are, and the timescale-scaling of its fluctuations.

In large-scale neural systems, the dynamics of individual brain regions have been shown to vary along a unimodal–transmodal axis [3], vary with structural connectivity strength [4, 5] and cortical microstructure [6], distinguish between individuals [7], characterize sleep states [8] and anesthesia [9], and be disrupted in autism spectrum disorder [10]. While many analyses of fMRI time series have focused on estimates of timescales—typically variations on the rate of decay of the autocorrelation function, or low-frequency spectral power—the existence of a large time-series feature library in *hctsa* allows us to compare a wide range of potential time-series properties. However, *hctsa* was designed as a comprehensive survey of the interdisciplinary time-series analysis literature and thus contains a large number of highly correlated features [1, 11].

For any given problem, or class of problems, we would like to distill the thousands of candidate features in *hctsa* down to a concise subset of the most informative time-series features that is efficient to compute and reduces the statistical comparison burden of the resulting analyses. Recent work demonstrated one approach to this problem: the thousands of *hctsa* features were reduced down to a high-performing and representative subset of 22 time-series features, called *catch22* [12]. This work defined high-performing features as those with the best performance across a diverse set of 93 time-series classification problems [13], and then used hierarchical clustering to reduce inter-feature redundancy, yielding a reduced set of 22 features. The 93 time-series classification tasks used to quantify ‘good’ performance spanned a wide range of general applications, from identifying birds using time series extracted from binary images [14] through to distinguishing types of coffee from their infrared spectra [15].

While *catch22* is a high-performing and efficient general-purpose time-series feature set, specific problems will require tailored features to optimally characterize the relevant dynamics [16]. Previous analyses of fMRI time series have involved two main issues. For studies that hand-select a single feature of interest (like a metric of autocorrelation timescale) [4, 5], the large range of alternative features are left untested, leaving open the possibility that alternative statistics could be more informative and interpretable. Applications that have systematically compared the performance across a large feature library, such as *hctsa* [3, 8, 17, 6, 18, 19], have involved substantial computational expense and statistical care in dealing with high-dimensional feature spaces (often requiring correction across thousands of features, limiting the power to detect signals in smaller samples). fMRI time series are short, noisy, and—particularly in the resting-state regime where dynamics may be approximated as close to a steady-state—are generally well-suited to being characterized by linear time-series analysis methods [20, 21]. However, some prior work has shown that novel time-series methods can reveal new insights into macroscopic brain organization: accurately tracking independent biological gradients in human brains [3], classifying stimulation experiments in mice [17, 19], and distinguishing experienced meditators from controls using resting-state EEG [18].

Here we aimed to develop a computationally efficient time-series feature set tailored to capturing informative dynamical patterns in fMRI time series, focusing mainly on the example of mouse fMRI (but later testing on independent datasets from both mouse and human). Successfully building an efficient reduced set of time-series features to characterize fMRI could thus facilitate new research into quantifying local dynamics in neuroimaging, allowing comparisons beyond conventional metrics, without the statistical and computational burden of evaluating thousands of candidate features. To judge the usefulness of different time-series features in this setting, we used data from recent experiments in mice using targeted chemogenetic manipulations of the underlying neural circuits (including firing of excitatory and inhibitory neurons) and measuring the fMRI dynamics in response [17]. These labeled data allowed us to score each time-series feature by how well it distinguishes biologically meaningful manipulations to the neural circuits. We introduce improvements to the *catch22* pipeline for constructing a reduced set of time-series features from a large feature library (such as *hctsa*) and use this improved pipeline to extract a set of 16 highly informative time-series features, which we term the *catchaMouse16* feature set and make it available as efficient open code for use by the community. We show that the performance of this set of 16 features in classifying different DREADD manipulations approximates that of the full set of > 7000 *hctsa* features and, consistent with it capturing biologically informative dynamical information, *catchaMouse16* also performs well on unseen tasks involving mouse fMRI.

## Methods

Our main aim was to reduce the comprehensive *hctsa* set of > 7000 candidate time-series features [2] down to a subset that displays similarly strong performance at distinguishing fMRI dynamics corresponding to biologically meaningful changes at the circuit-level, assessed using a dataset of chemogenetic manipulations to mouse cortical circuits [17]. In this section, we describe the data and corresponding classification tasks used to score the performance of individual time-series features. We then describe our method for setting the parameters of a feature-reduction pipeline, adapted from Lubba et al. [12], using a leave-one-task-out cross-validation approach. Code to reproduce our analysis is available at https://github.com/DynamicsAndNeuralSystems/catchaMouse16_paper_code.

### Dataset

Here we consider a set time-series classification tasks based on distinguishing different chemogenetic manipulations from fMRI time series. We use data described in Markicevic et al. [17], where full experimental details can be found; key information is summarized here. Chemogenetic manipulations were made in the primary somatosensory area (SSp) in the right hemisphere of the mouse isocortex. As well as the control condition (labeled ‘SHAM’ here), three chemogenetic manipulations were performed: (i) activating CaMKIIa-hSyn-hM3Dq-DREADD in wild-type mice, which increases overall neural excitability (labeled ‘excitatory’ here); (ii) activating hM3Dq-DREADD in wild-type mice, which selectively increases the excitability of pyramidal neurons (labeled ‘CAMK’ here); and (iii) activating inhibitory hSyn-dlox-hM4Di-DREADD in PVCre mice, which selectively decreases the excitability of PV interneurons (labeled ‘PVCre’ here).

From this dataset, we constructed a total of 12 classification tasks: six based on data from the right-hemisphere SSp (spatial site of the injected DREADD) and six based on data from the contralateral left-hemisphere SSp. In each region, the six classification tasks corresponded to all pairwise combinations of the four labeled classes: ‘SHAM’ (13 fMRI time series), ‘excitatory’ (14), ‘CAMK’ (13), and ‘PVCre’ (19). We labeled each of these twelve tasks in the form `region-group1_group2`, such that `r-SSp-CAMK_PVCre` corresponds to the task in which the goal is to distinguish CAMK and PVCre manipulations in the right-hemisphere SSp.

### Feature extraction, scoring, and reduction

We used *hctsa* (v1.03) to extract the full *hctsa* feature set from all 118 (59 left-hemisphere and 59 right) time series in the dataset. Missing or special-valued outputs that occurred in any of the 12 classification tasks were ignored in subsequent calculations. After filtering, all remaining features were normalized as a *z*-score. A total of *F* = 6356 *hctsa* features survived filtering across all tasks and were included in the remaining analysis. In each task, every feature was scored according to its balanced classification accuracy using a linear support vector machine (SVM) using the `SVM` module of `scikit-learn` [22]. The results were then represented as a task × feature (12 × *F*) performance matrix (where each element in the matrix is a measure of the performance of a given feature in a given task).

From the performance results per feature, we aimed to determine a reduced set of features that exhibit strong performance across the set of classification tasks while minimizing inter-feature redundancy, as per Lubba et al. [12]. We follow a similar two-stage approach as Lubba et al. [12], first selecting the *β* features with the highest average balanced classification accuracy across the 12 tasks, and then clustering them to reduce redundancy. For this second step, we used hierarchical average linkage clustering, replacing each cluster of highly correlated features with a single representative feature. We defined feature–feature dissimilarity as 1 − |*R*|, where *R* is the Pearson correlation between their performance values across the 12 classification tasks [12]. In this way, groups of features with similar patterns of classification strengths and weaknesses across classification tasks cluster together (and are therefore considered ‘redundant’ with each other). We cut the resulting dendrogram at a threshold *γ* and represented each cluster as the highest-performing feature in the cluster where possible and, when performance differences were minimal, used scientific judgement when replacing some computationally expensive features with simpler and more interpretable alternatives within a cluster to enhance speed and interpretability. In Lubba et al. [12], *γ* was set manually to *γ* = 0.2, corresponding to clusters containing features with highly correlated performance |*R*| > 0.8. We used ‘average’ linkage clustering here as we found that it yields superior cross-validated results than the stricter ‘complete’ linkage clustering used previously [12] (see Fig. S2).

This pipeline for constructing reduced sets of features thus depends on the selection of *β* and *γ*. Here we extend the approach of Lubba et al. [12] by introducing a data-driven approach to select the values of *β* and *γ*, using leave-one-task-out cross-validation. In this scheme, for a given setting (*β, γ*), a reduced feature set was constructed using data from 11 of the 12 datasets, leaving the left-out dataset as an out-of-sample test. Performance in the left-out dataset was taken to be the 10-fold cross-validation balanced accuracy, using the feature set constructed from the 11 training datasets. The performance of a reduced time-series feature set obtained using given parameters, (*β, γ*), was calculated as the average classification performance of that feature set across all 12 left-out tasks. This procedure allowed us to select a value for *β* and *γ* to then use on the full set of 12 datasets to produce a final set of features, *catchaMouse16*.

## Results

Our results are organized as follows. First, we describe our leave-one-task-out method for selecting appropriate parameters of our feature-reduction pipeline and how this yields the *catchaMouse16* feature set. We then characterize the composition of this feature set and evaluate its performance relative to the entire *hctsa* feature set. Finally we show that *catchaMouse16* performs well in independent datasets of mouse and human fMRI time series, in which cortical variations in cell densities are well captured by the variation of *catchaMouse16* features of the resting-state fMRI dynamics.

### Computing a reduced feature set

In this section, we describe our leave-one-task-out procedure for generating the reduced feature set, *catchaMouse16*, from the twelve classification tasks derived from Markicevic et al. [17]. To understand how the clustering threshold *γ* affects performance on left-out tasks, we fixed *β* = 100 and explored the accuracy on each of the twelve tasks across a set of thresholds *γ* = 0.05, 0.1, 0.2, 0.3, 0.4, 0.5. Note that for each left-out dataset, the 11 training datasets are used to select a reduced set of features, and a classification model is then trained and evaluated on the left-out dataset.

Results are shown for all tasks and thresholds in Fig. 1A, showing the number of features selected from the other tasks (numerical annotations) and balanced accuracy on that left-out task (color). We first note that clustering at higher thresholds, *γ*, corresponds to larger clusters and thus fewer features in the final feature set, with performance broadly degrading at high thresholds, *γ* (corresponding to fewer features). The trade-off involved in the feature-set reduction problem is thus clearly visible as aiming to produce as high out-of-sample performance while using as few features as possible.

**Figure 1.**
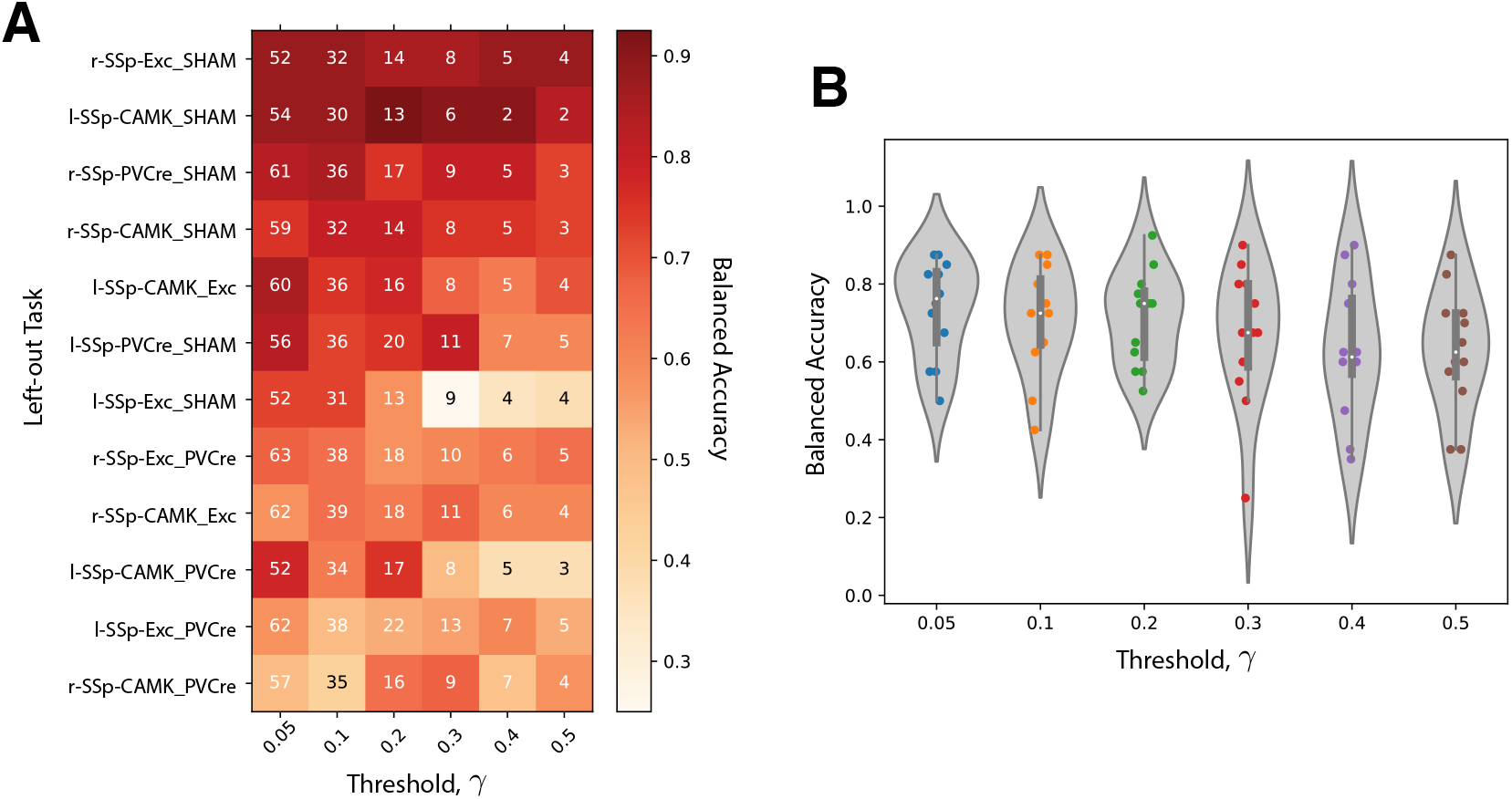
A leave-one-task-out procedure allowed us to select a high-performing clustering threshold, *γ*. Results are shown here for the example of clustering the *β* = 100 highest-performing features across the 12 tasks down to a reduced subset, comparing the results of using *γ* = 0.05, 0.1, 0.2, 0.3, 0.4, 0.5. **A** We plot this task × threshold matrix that shows the results of the leave-one-task-out procedure as a heat map. The number of features used in the model (selected from the other tasks) is annotated to each element, and the balanced accuracy in classifying the left-out-task is shown as color. Left-out tasks (rows) have been sorted by mean balanced accuracy from high (top) to low (bottom). **B** The distribution of balanced accuracies aggregated across all tasks (i.e., aggregating down columns of the matrix shown in A is shown as a violin plot for each threshold *γ*. Note that the number of retained features decreases with *γ*, and varies across left-out tasks, as annotated in A.

Comparing tasks (rows) of Fig. 1A, we see a range of accuracies across the different tasks: high performance is obtained on some tasks (even with few features), while other tasks display relatively poor performance. The four tasks with the highest classification accuracy in the original analysis by Markicevic et al. [17]—`r-SSp-Exc_SHAM`, `l-SSp-CAMK_SHAM`, `r-SSp-PVCre_SHAM`, and `r-SSp-CAMK_SHAM`—are also the four tasks with highest performance here. High performance on these tasks was mostly maintained across *γ*, even at high *γ* = 0.5 we see similar classification rates with just a few features (selected from the other 11 tasks). By contrast, tasks like `l-SSp-Exc_SHAM` and `l-SSp-CAMK_PVCre`, exhibit a substantial drop in performance when going from a large feature set (low *γ*) to a smaller feature set (high *γ*).

To select a value for *γ* to apply to the full dataset in constructing *catchaMouse16*, we examined the distributions of left-out performance, shown in Fig. 1B. The median accuracy across left-out tasks remains approximately constant for *γ* = 0.05, 0.1, 0.2 but drops from *γ* = 0.3. This suggests that we can maintain performance while reducing the size of the feature set by selecting *γ* = 0.2.

Having selected parameters *β* = 100, *γ* = 0.2, we applied our feature-reduction pipeline with these settings to the full set of 12 classification problems to compute a reduced time-series feature set. This yielded a set of 16 clusters of high-performing features, shown as an annotated feature–feature correlation matrix in Fig. 2. We then selected a representative feature from each cluster, using scientific judgement to select a high-performing representative that was as simple to interpret and computationally inexpensive as possible, as per Lubba et al. [12]. The resulting set of 16 features is the *catchaMouse16* set.

**Figure 2.**
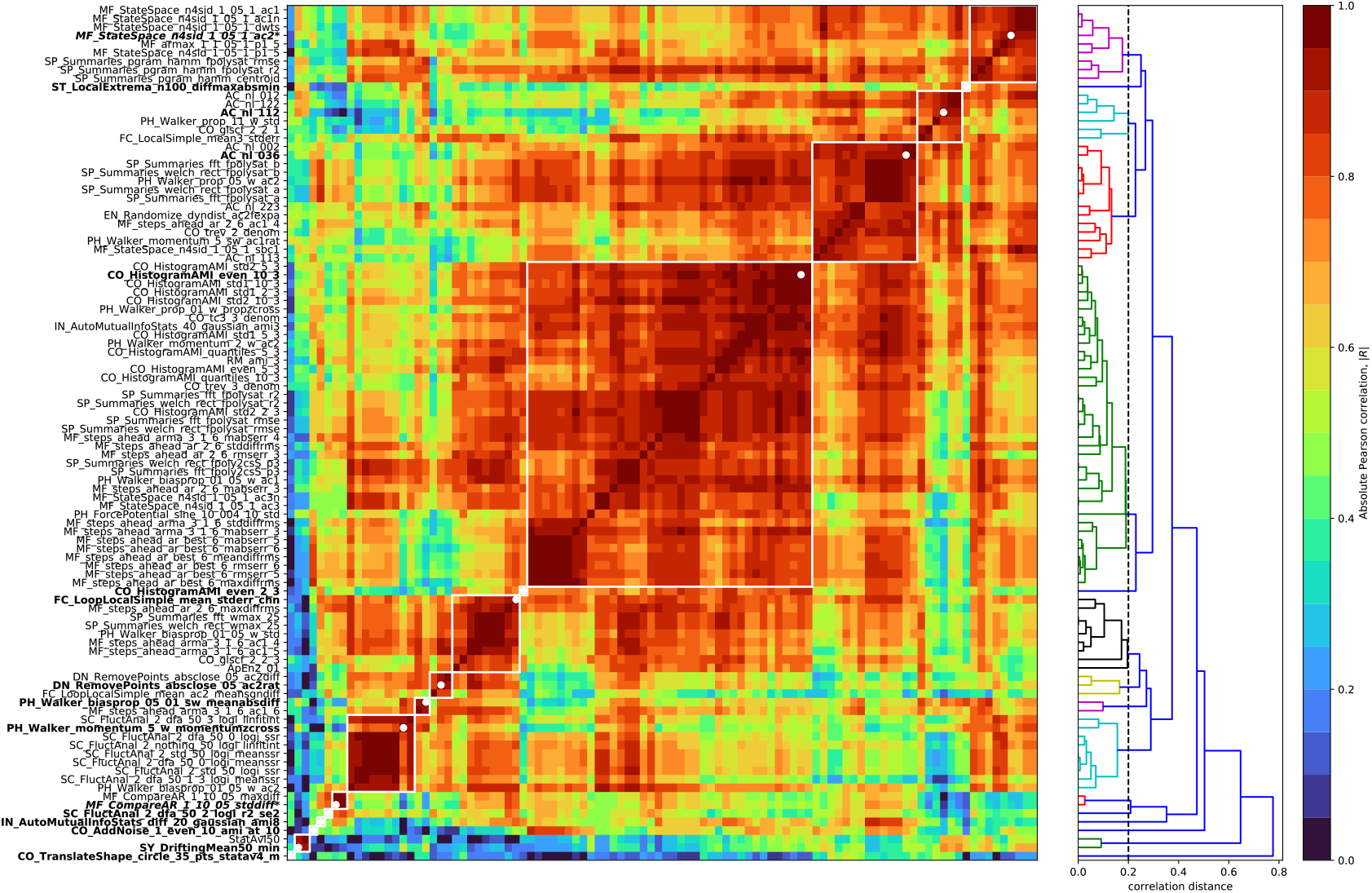
Feature–feature correlation matrix of the 100 top-performing features across the 12 mouse fMRI classification tasks. A heatmap shows the correlations between each pair of features, as |*R*|, computed from the variation in performance values across the 12 tasks. Features in the *catchaMouse16* feature set are denoted in bold, while the italic features marked with an asterisk denote features which were substituted for similar but easier-to-compute features. The average hierarchical linkage dendrogram used to determine the clusters at a threshold *γ* = 0.2 (annotated) is shown to the right of the heatmap.

### The *catchaMouse16* feature set

Having determined a reduced set of 16 time-series features, designed to be concentrated in the types of dynamical structures that are most relevant to changes in the underlying neural circuits, we next describe the time-series features in the *catchaMouse16* feature set. The features are listed in Table S1, where we have assigned a short name to each feature for ease of use. The *catchaMouse16* features mainly comprise measures of predictability, such as nonlinear autocorrelations, automutual information estimates, and estimates of long-range scaling of correlation structure.

Three *catchaMouse16* features compute different nonlinear time-averages of the *z*-scored time series *x*_*t*_, as: 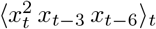 (`nonlin_autocorr_036`), 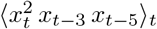 (`nonlin_autocorr_035`), and 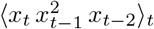 (`nonlin_autocorr_112`). Two other features measure nonlinear time-lagged correlations via automutual information: `ami3_10bin` and `ami3_2bin` measure the automutual information at a time-lag of 3 samples using a 10-bin (and 2-bin) histogram estimation method, respectively. Another feature, `increment_ami8`, measures linear time-lagged correlation at a lag of 8 samples, using a linear Gaussian estimator for automutual information.

A range of other *catchaMouse16* features capture different aspects of time-series predictability. The `noise_titration` feature is inspired by the noise titration method of Poon and Barahona [23], which measures the lag-1 automutual information after adding white measurement noise (with a signal-to-noise-ratio of 1) to the time series. And `prediction_scale` captures the change in prediction error from a mean forecasting method when incorporating increasing durations of prior data into the prediction. The `dfa_longscale_fit` feature estimates a timescale–fluctuation curve using detrended fluctuation analysis (DFA) [24] using quadratic detrending in each window (and returns the standard error in the mean of a robust fit to a long-timescale fluctuation scaling regime). Other more novel features of local shape-like patterns in the time series include the variability of how many time-series points sit inside a circle being translated across the time domain (`floating_circle`), and two measures of the statistics of a simulated mechanical particle (or ‘walker’) being driven by the time series (`walker_crossings` and `walker_diff`) [1, 2].

Two *catchaMouse16* features capture aspects of the stationarity of the signal [16]. First, `stationarity_min` divides the time series into 50 windows and then measures the minimum windowed mean in any segment divided by the mean variance in windows. And `stationarity_floating_circle` measures the stationarity across four partitions of the time series (using a quantity called StatAv [25]) of the statistics of local time-series shapes (number of points sitting inside circles translated across the time series).

Finally, *catchaMouse16* contains two measures related to outliers in the time series. `outlier_corr` computes how the autocorrelation at lag 2 changes after removing 50% of the time-series values closest to the mean. And `outlier_asymmetry` measures asymmetry in extreme local events by dividing the time series into 100 segments and computing the average difference between the greatest deviation in positive and negative directions.

Together, the *catchaMouse16* features aim to provide a concise but multifaceted signature of BOLD dynamics, covering linear and nonlinear correlation structure, scaling behavior, statistics of local shapes, stationarity, and outliers. For broader use, this feature set has been re-coded from the original Matlab into C, and is made available as an open-source package, *catchaMouse16*, with wrappers for Matlab and Python, available at https://github.com/DynamicsAndNeuralSystems/catchaMouse16.

We analyzed the performance of the *catchaMouse16* feature set on the 12 classification tasks from Markicevic et al. [17]. As shown in Fig. 3A, the *catchaMouse16* features yield similar performance as the full *hctsa* feature set across the 12 tasks. Further, the feature set performs similarly to the centroids of the feature clusters as outlined in Fig. 2, with the added advantage of being more clearly interpretable and simpler to compute (see supplementary Fig. S1 for details). Although this analysis is ‘in-sample’ in the sense that it is performed on the same dataset on which the *catchaMouse16* features were constructed (analyses on independent datasets are presented in the next section), these results indicate that *catchaMouse16* can differentiate underlying neural stimulations from non-invasively measured fMRI time series using just 16 features (with similar or greater accuracy as the full *hctsa* set of ∼ 7000 features).

**Figure 3.**
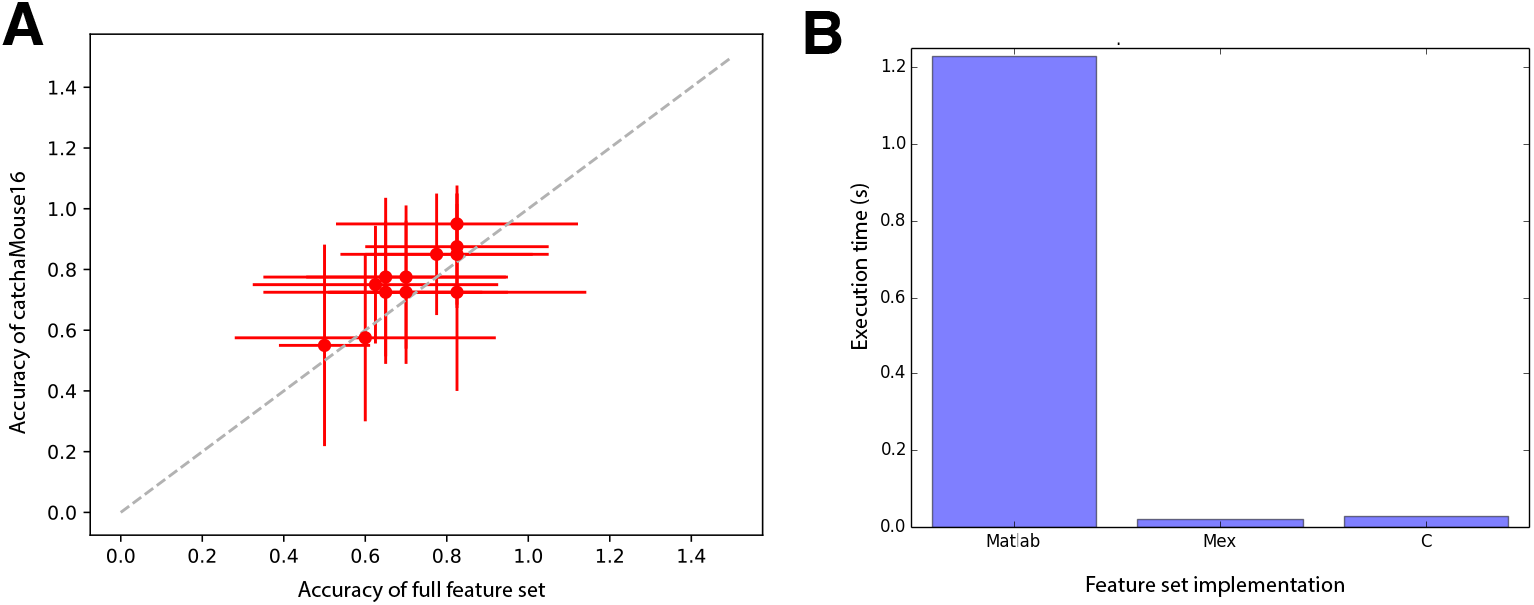
The *catchaMouse16* feature set maintains high performance with substantial gains in computational efficiency relative to *hctsa*. **A**: Mean 10-fold cross validated balanced classification accuracy of *catchaMouse16* (vertical axis) versus *hctsa* (horizontal axis) across the twelve classification tasks analyzed here. Error bars indicate standard deviations across folds. **B**: Execution time of the *catchaMouse16* feature set in native Matlab code, C code run through Matlab (mex), and in compiled C.

To characterize the relative computational efficiency of the Matlab implementation of the *catchaMouse16* feature set compared to the C-coded version introduced here, we evaluated the feature set on i.i.d. random-number time series of length 900 (repeating the process 2000 times). Results are shown in Fig. 3B, and indicate a ∼ 60× speed-up of the C implementation (and Matlab mex wrappers of it) relative to the native Matlab code from *hctsa*. This makes *catchaMouse16* highly efficient at quantifying dynamical structure in time series that makes it suitable for scaling up to large time-series datasets.

### Validation: tracking cell-density variation across brain areas

In this section, we aimed to test the generalization ability of *catchaMouse16* (which aims to be an efficient and information-concentrated subset of *hctsa*) on new problems involving fMRI data. Specifically, we aimed to detect statistics of resting-state BOLD dynamics that vary with the relative density of different neuronal types across brain areas (in both mouse and human). In mouse, we focused on pyramidal (excitatory) neurons as well as the three main subtypes of inhibitory interneurons: parvalbumin (PV), vasoactive intestinal peptide (VIP), and somatostatin (SST) expressing interneurons [26]. While the densities of all classes of interneurons vary substantially across the cortex [27, 28], the precise roles of interneuron subtypes in shaping macroscopic neural activity remain unclear [29, 30]. Identifying time-series features of fMRI dynamics that correlate strongly with the density of interneuron subtypes would be a step toward connecting non-invasive measurements of macroscopic dynamics with the microscopic properties of neuronal circuits. We compared the performance of both *hctsa* and *catchaMouse16* time-series features: if the smaller *catchaMouse16* feature set can capture the main variation as well as the full *hctsa* feature set, then this would support it being an information-rich set of informative dynamical properties of fMRI time series.

We used a dataset of resting-state fMRI recordings from 38 regions of the isocortex in the right hemisphere of 100 mice (each of length *T* = 900 samples) [31, 32, 5] (parcellated according to the Allen Mouse Brain atlas [33]). For each feature-extraction pipeline (either *hctsa* v1.03, 7664 time-series features; or *catchaMouse16*, 16 time-series features), we extracted features from each of the 3800 BOLD time series. We then agglomerated the result for each feature and each brain region by calculating a mean value across all 100 mice (excluding any missing or `NaN` values). In the case of the *hctsa* feature set, this yielded a 169 × 7664 averaged matrix (region × features).

In mouse, we compared the spatial variation of each time-series feature (across brain areas) to data on their relative cellular composition obtained from two sources: (i) densities of PV, VIP, and SST interneurons, obtained by cell counting [27]; and (ii) densities of excitatory and inhibitory populations, estimated from an algorithmic reconstruction of neuronal soma [34]. We combined these two sources of cell-density measurements to produce a table of PV, SST, VIP, and inhibitory cell densities for each region in the mouse brain. We repeated the same approach on human resting-state fMRI data obtained from 100 participants in the Human Connectome Project [35]. These data, comprising processed and parcellated time series for 180 left-hemisphere brain regions using the Glasser parcellation [36], were taken from Fallon et al. [4]). As a proxy for PV interneuron density in the human cortex, we used expression of the *PVALB* gene, taken from Arnatkeviciūtė et al. [37]. We chose to focus just on the *PVALB* gene in humans as it is the most reliable marker for the corresponding interneuron type (PV) in mouse (as found here, cf. Fig. 5, and in prior work [38]), whereas the correspondence between genetic markers for SSP and VIP and their ground-truth densities is less clear [38]. In both the mouse and human datasets, we quantified the relationship between neuron densities and statistical features of BOLD time series across brain areas using the Spearman correlation, *ρ*. We assessed corrected statistical significance using the Bonferroni–Holm method [39], noting that such correction is much more harsh for the *hctsa* feature set (where thousands of features are tested) than the *catchaMouse16* feature set (where only 16 features are tested).

We first analyzed the relationship between fMRI time-series features and cell densities in the 38 regions of the mouse isocortex and *PVALB* expression in the 180 regions of the human cortex. Of the six cell-type density associations tested, four exhibited a significant correlated with at least one time-series feature: PV, VIP, and inhibitory cells in mouse, and *PVALB* expression in human. In the other two cases, excitatory cell density and SST cell density in mouse, there were no significant correlations to any time-series features of BOLD dynamics (in neither *hctsa* nor *catchaMouse16*). Figure 4 shows the distributions of squared correlations, *ρ*^2^, across all BOLD time-series features in *catchaMouse16* and *hctsa* for the four settings in which there was at least one feature with a significant correlation. The distributions of *hctsa* features are shown as violin plots, with *catchaMouse16* features overlayed as circular markers and features with significant correlations (*p*_corr_ < 0.05) highlighted in color. For *hctsa*, we observed significant correlations between individual time-series features of resting-state BOLD dynamics and mouse PV cell density (240 significant features, up to *ρ* = 0.75, *p* < 10^−3^), mouse VIP cell density (1 significant feature: lag-21 autocorrelation, *ρ* = −0.70, *p* = 7 × 10^−3^), overall mouse inhibitory cell density (249 significant features, up to *ρ* = 0.78, *p* < 10^−4^), and human *PVALB* expression (273 significant features, up to *ρ* = 0.32, *p* = 2 × 10^−4^). A complete list of correlations and values for each *hctsa* feature and each cell type is given in Supplementary files `PV_7664.csv`, `VIP_7764.csv`, and `Inhibitory_7764.csv`, and, for human, `Pvalb_7764.csv`.

**Figure 4.**
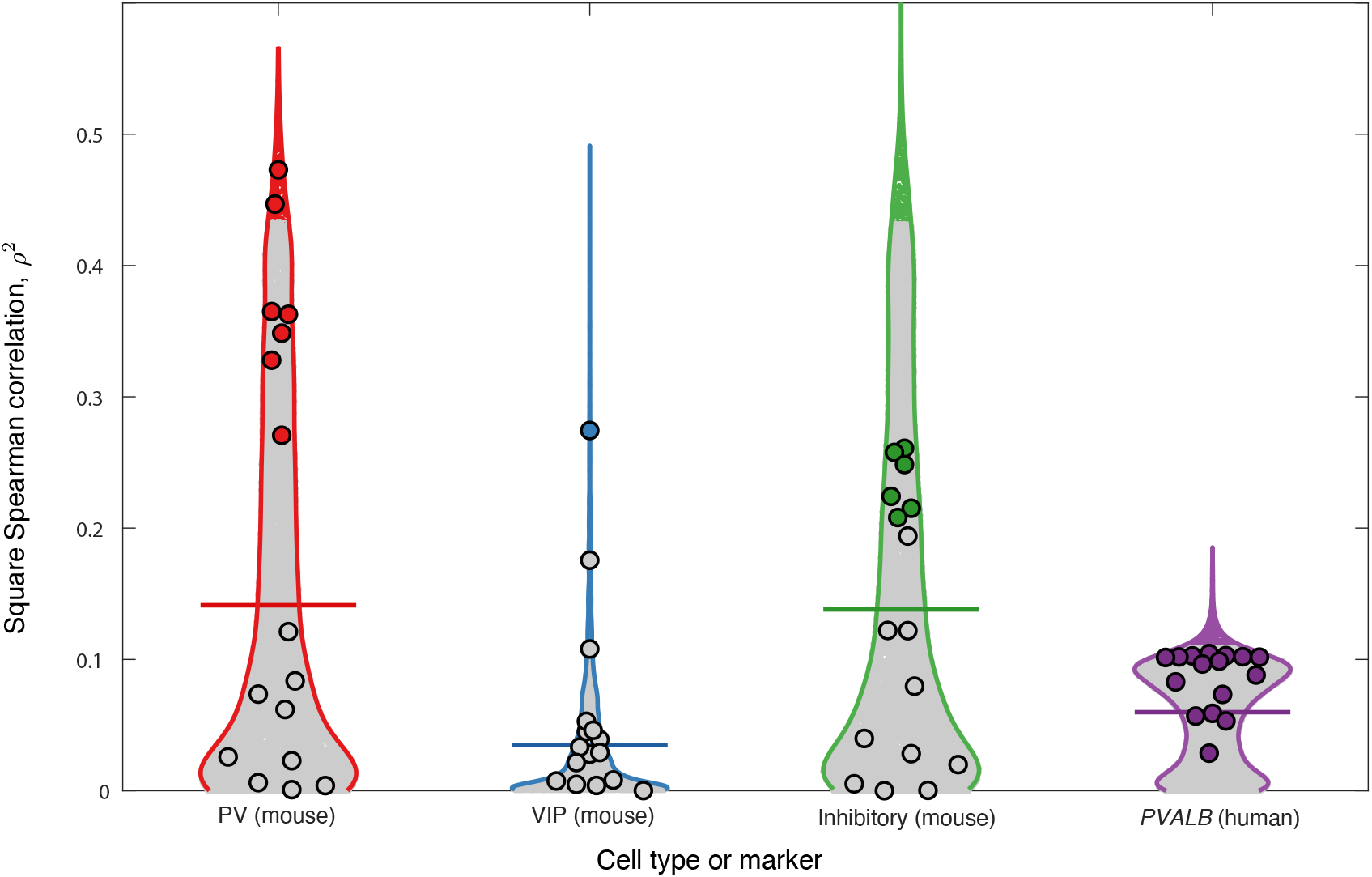
*catchaMouse16* efficiently captures dynamical properties of mouse and human resting-state BOLD time series that are informative of regional variations in interneuron density across the cortex. The distribution of squared Spearman correlation coefficients, *ρ*^2^, between *hctsa* feature values of BOLD time series and estimated interneuron cortical cell densities are shown as a violin plot for each cell type or marker. Gray points represent features with *p*_corr_ ≥ 0.05; colored points represent features with *p*_corr_ < 0.05. The mean correlation across *hctsa* features is indicated by a horizontal line. The 16 *catchaMouse16* features are annotated as larger circles, shown as filled gray circles (*p*_corr_ ≥ 0.05) or filled colored circles (*p*_corr_ < 0.05). As annotated, the PV, VIP, and inhibitory cell densities correspond to mouse data, whereas the *PVALB* marker is from human data.

Using the *catchaMouse16* feature set yielded a significant relationship in all settings that *hctsa* did. Furthermore, the proportion of *catchaMouse16* features that were significantly correlated with each anatomical measurement was far greater than for *hctsa*, with the top-performing *catchaMouse16* feature mostly yielding a similarly strong correlation to the top *hctsa* feature (in all cases except the overall inhibitory density analysis). The strongest associations to *catchaMouse16* features also varied in interesting ways between the cell types. For example, the *catchaMouse16* feature with the strongest correlation to the spatial variation in PV cell density was `nonlin_autocorr_036` (*ρ* = −0.69, *p*_corr_ < 10^−4^, cf. Table S2 for all results), with similar types of self-correlation-based time-series features performing well at tracking total inhibitory cell density, including `nonlin_autocorr_036` (*ρ* = 0.5, *p*_corr_ = 0.02), although with weaker *ρ*^2^ than the top *hctsa* features. By contrast, VIP density exhibited the strongest correlation to `dfa_longscale_fit` (*ρ* = 0.52, *p*_corr_ = 0.01, cf. Table S3 for all results). Overall, the features with the highest *ρ* with respect to the inhibitory cell density had lower *ρ* with respect to the VIP cell density; for example, `noise_titration` has *ρ* = −0.51 (*p*_corr_ = 0.02) for inhibitory cells, and *ρ* = 0.086 (*p*_corr_ = 1) for VIP cells (cf. Table S4 and Table S3). In the human cortical *PVALB* analysis, all *catchaMouse16* features exhibited a significant correlation (with the strongest correlations being with features capturing nonlinear autocorrelations, like `nonlin_autocorr_036`, *ρ* = 0.32, *p* < 10^−3^, and automutual information, like `ami3_2bin`, *ρ* = 0.32, *p* < 10^−3^, cf. Table S5). These results suggest that *catchaMouse16* can act as a compact and computationally efficient distillation of the time-series analysis literature that is suitable for capturing biologically relevant properties of fMRI dynamics.

### Layer-specific study

As a final study of the statistical relationship between interneuron densities and fMRI dynamics in mouse cortex, we broadened our scope to compare the *catchaMouse16* time-series features of the regional BOLD signal with layer-specific interneuron densities. Figure 5 shows the maximum correlation (annotated as text) of *hctsa* and *catchaMouse16* features to the density of each interneuron subtype in the isorcotex as well as each layer of the isocortex (5 layers over 38 regions). This figure also annotates the number of significant features in each feature set (as a heatmap) for each layer and cell type. Comparing Figs 5A and B shows that *catchaMouse16* is able to identify features that are significantly correlated with interneuron densities at a much higher rate than *hctsa*, due to the concentration of informative time-series features and the much lower correction burden of multiple hypothesis testing. And, for cell type and layer pairs that have many significant features, the top *catchaMouse16* correlations are comparable to the top *hctsa* correlations. Although these results come from performing many comparisons on a single dataset, we note that some *catchaMouse16* features track some layer-specific cell densities with a striking level of correspondence, e.g., `ami3_10bin` tracks inter-areal variation in layer 4 VIP densities with *ρ* = −0.93. Such patterns suggest promising areas for future work to investigate further.

**Figure 5.**
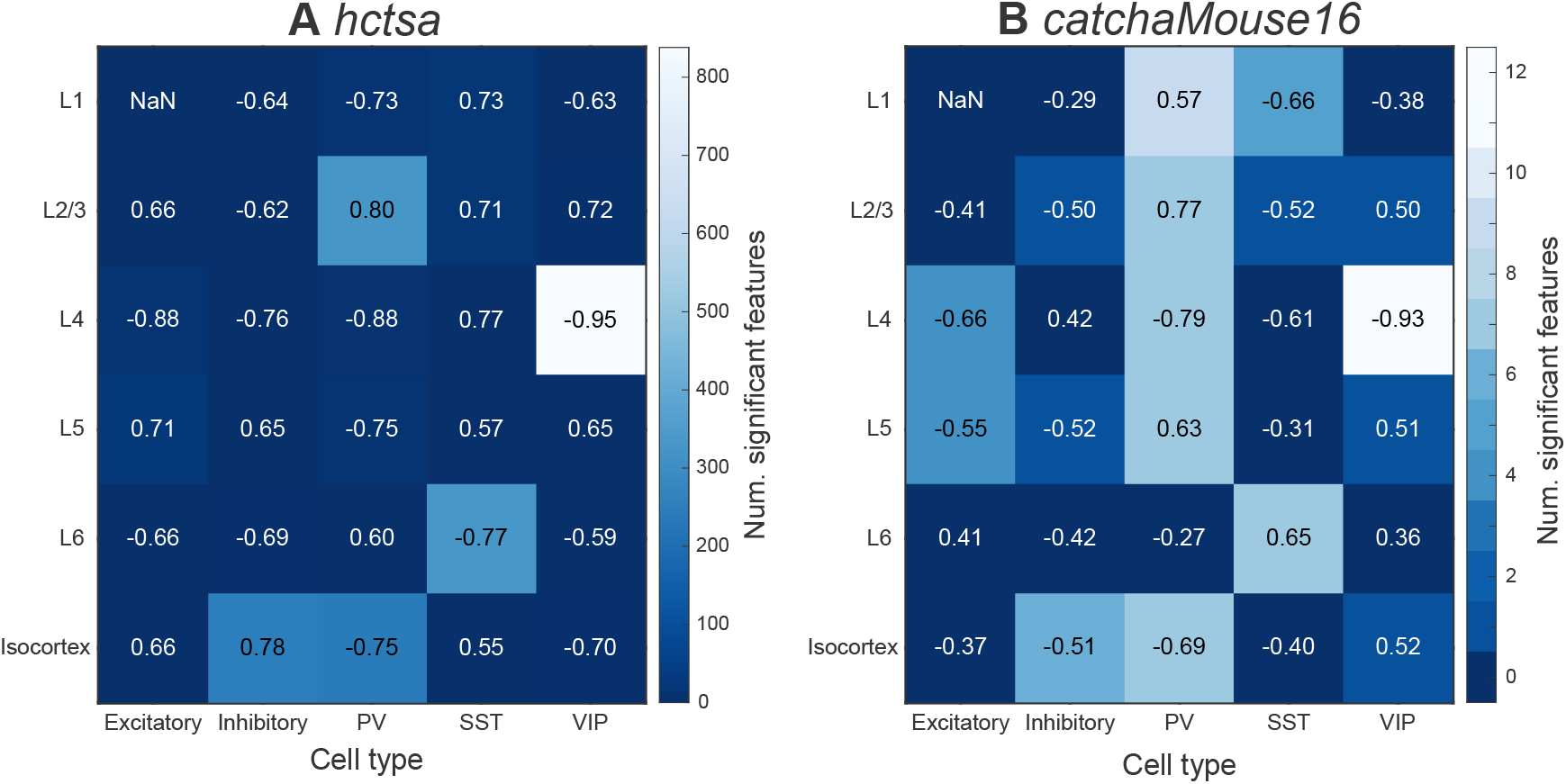
*catchaMouse16* is enriched in features that are significantly correlated to the cortical variation of interneuron cell densities in all cytoarchitectural layers of the mouse cortex. The number of features with significant correlations to regional neuronal densities (*p*_corr_ < 0.05, Bonferroni–Holm corrected across all features within a given feature set), is shown in a heatmap for both *hctsa* (**A**) and *catchaMouse16* (**B**). Each annotated number corresponds to the maximum Spearman’s correlation for a combination of layer and cell type. Rows 1-5 were calculated using cell-type densities in one of layers 1 − 6 in the mouse isocortex, while row 6 uses densities across all layers in each isocortical region. Note the color axis of **A** spans approximately 10% of the *hctsa* features, whereas the color axis of **B** spans most of the *catchaMouse16* features.

## Discussion

In this work we aimed to reduce the comprehensive but computationally expensive *hctsa* feature library down to an efficient and minimally redundant subset that is enriched in time-series properties relevant for tracking biologically meaningful variation from the BOLD signal. We adapted an existing pipeline from Lubba et al. [12] to achieve this, assessing feature performance based on a set of brain stimulation experiments from Markicevic et al. [17]. The resulting set of 16 interpretable features, *catchaMouse16*, is vastly more efficient to compute than *hctsa*, and mitigates the multiple hypothesis ambiguity associated with comparing thousands of time-series features. To demonstrate its utility, we present a novel analysis on an independent dataset containing variations in cell-density across mouse and human cortical areas, where we demonstrate that the *catchaMouse16* feature set is indeed enriched in relevant dynamical properties of the fMRI signal (relative to *hctsa*), with its smaller size sometimes resulting in greater levels of statistical significance (due to a smaller multiple hypothesis correction burden than *hctsa*). Our results also provide interpretable insights into the types of properties of non-invasive fMRI measurements that can track underlying cell-type variations in neural circuits, an important precursor to using non-invasive imaging as a kind of ‘cellular microscope’. Indeed, our results speak to an exciting potential of cross-species generalizability, with similar types of time-series features showing strong correlations to PV density across mouse cortical areas as to the relevant gene marker (*PVALB* expression) in human.

The methodology demonstrated here provides a path to producing efficiently coded reduced time-series feature sets for a range of applications and settings and improves on that developed by Lubba et al. [12] in important ways. Relative to the Lubba et al. [12], we introduce a leave-one-task-out design, and provide new ways of selecting hyperparameters such as the clustering method and threshold *γ*. There is room for further improvements in future work, by explicitly incorporating feature-selection methods, that go beyond scoring features individually and take into account synergistic performance of combinations of features. For example, such extensions could build from simple greedy forward-selection methods [40]. While our method has the advantage that every feature in the reduced feature set is individually a high-performing feature, a feature-selection approach is likely to yield improved performance with smaller feature sets in multivariate settings. We also highlight the limitation of using a single set of experiments to judge ‘high-performing’ features; A clear direction for future work is to incorporate a larger database of problems in the training of the reduced feature set, such as correlations with other anatomical features (as treated in the cell-type correlation results here) [41], and the myriad of other fMRI classification tasks, from case–control, gene-knockout–wildtype, to task–rest comparisons (e.g., cf. [42]).

Recent work has highlighted the benefits of using sophisticated univariate time-series analysis methods to analyze neural dynamics, even for modalities like fMRI, where relatively high noise levels and low temporal resolution mean that linear methods are expected to perform well [42, 3, 6, 4, 20, 21]. Here, many novel dynamical properties (and corresponding parts of the time-series analysis literature) that are not traditionally used in fMRI time-series analysis—like non-linear autocorrelations (such as 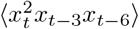, measures of stationarity, and outlier properties— are found here to be high-performing candidate features for fMRI analysis. Interestingly, the *catchaMouse16* feature set does not include classical linear time-series analysis methods, like statistical properties of the (linear) autocorrelation function or power spectrum. Although our results here are based on a single set of experiments, and the feature set presented here is accordingly not definitive, these results suggest that there may indeed be some promise in applying diverse types of time-series analysis methods to short, noisy fMRI time series [3].

Our validation case studies, tracking the variation in cell density (and cell-density markers) across cortical areas in mouse and human brains, flag biologically interesting results in their own right, showing that gradients of cortical organization, resolved at the level of individual cortical layers, can be tracked by relevant univariate statistics of resting-state BOLD dynamics [43, 38, 5]. These findings provide new evidence for the relationship between the cellular composition of neural circuits and the macroscopic dynamics of large-scale neural populations [29, 44, 43, 45, 46]. Some particularly strong correlations provide particular directions for future investigation, such as our finding that layer 4 VIP cell density correlates strongly (up to |*ρ*| = 0.95) with specific dynamical properties of resting-state BOLD dynamics. While such strong correlations await additional experimental verification on independent datasets, our results point to promising candidate layer-specific and cell-type specific mechanisms that may strongly shape the emergent population-level dynamics. This is important because population-level activity can be detected non-invasively, and may provide a window into the underlying neural circuits in mouse, and their correspondence in human [43].

While the unification of the interdisciplinary time-series analysis literature in *hctsa* has overcome some of the barriers of applying diverse methods to real-world applications, it was designed for its comprehensive coverage of methods and, as such, contains large amounts of redundancy. The tailored reduction to *catchaMouse16* introduced here overcomes key barriers to using *hctsa* (including its computational expense and requirement of a proprietary Matlab software license), and has additional benefits of being a compact and interpretable set of features that can enable efficient analyses of dynamic structure in fMRI time-series data at scale. This includes the ability to run univariate time-series analysis on a finer spatial scale (e.g., voxel-wise feature extraction) than would be tractable using *hctsa*, and to run feature extraction across large databases of fMRI time series [32]. And although *catchaMouse16* was trained here on mouse fMRI (where we have a clear signal from direct biological manipulations [17]), and is named accordingly, our case study provides some preliminary evidence of *catchaMouse16* also being a useful tool for characterizing dynamical structure in human fMRI. We hope that the pipeline demonstrated here for constructing reduced feature sets for specific applications can yield more precisely tailored sets for other modalities and settings in future, by incorporating a greater range and diversity of tasks (e.g., calcium imaging, human MEG, etc.). Further, the open source *catchaMouse16* feature set that accompanies this paper (with the ability to be called in multiple coding languages) provides an accessible way for researchers to quantify useful dynamical structure from time-series data, potentially directing them to explore novel and fruitful areas of the interdisciplinary time-series analysis literature that are relevant for a given analysis problem.

## Acknowledgments

BDF would like to thank the Selby Scientific Foundation for financial support. VZ is supported by the Swiss National Science Foundation (SNSF) ECCELLENZA (PCEFP3_203005). The authors would like to thank Annie Bryant for helpful comments on the manuscript.

