## Supplementary Tables and Figures for "Canonical time-series features for characterizing biologically informative dynamical patterns in fMRI"

July 15, 2024

| Feature Name | *hctsa* Feature Name | Description |
|---|---|---|
| nonlin_autocorr_035 | AC_nl_035 | $\langle x_t^2\, x_{t-3}\, x_{t-6}\rangle_t$ |
| nonlin_autocorr_036 | AC_nl_036 | $\langle x_t^2\, x_{t-3}\, x_{t-5}\rangle_t$ |
| nonlin_autocorr_112 | AC_nl_112 | $\langle x_t\, x_{t-1}^2\, x_{t-2}\rangle_t$ |
| ami3_10bin | CO_HistogramAMI_even_10bin_ami3 | Automutual information at lag 3 using a 10-bin histogram estimation method. |
| ami3_2bin | CO_HistogramAMI_even_2bin_ami3 | Automutual information at lag 3 using a 2-bin histogram estimation method. |
| increment_ami8 | IN_AutoMutualInfoStats_diff_20_gaussian_ami8 | Mutual information at time lag 8 using Gaussian estimator. |
| dfa_longscale_fit | SC_FluctAnal_2_dfa_50_2_logi_r2_se2 | Timescale–fluctuation curve using DFA. |
| noise_titration | CO_AddNoise_1_even_10_ami_at_10 | Automutual information at lag 1 after adding white noise at a SNR of 1. |
| prediction_scale | FC_LoopLocalSimple_mean_stderr_chn | Change in prediction error from a mean forecaster using prior windows of data. |
| floating_circle | CO_TranslateShape_circle_35_pts_std | Variability of time-series points inside a circle translated across the time domain. |
| walker_crossings | PH_Walker_momentum_5_w_momentumzcross | Statistics of a simulated mechanical particle driven by the time series. |
| walker_diff | PH_Walker_biasprop_05_01_sw_meanabsdiff | Statistics of a simulated mechanical particle driven by the time series. |
| stationarity_min | SY_DriftingMean50_min | Minimum mean across 50 segments divided by mean variance in segments. |
| stationarity_floating_circle | CO_TranslateShape_circle_35_pts_statav4_m | StatAv of the statistics of local time-series shapes. |
| outlier_corr | DN_RemovePoints_absclose_05_ac2rat | Change in lag-2 autocorrelation from removing 50% of the time-series values closest to the mean. |
| outlier_asymmetry | ST_LocalExtrema_n100_diffmaxabsmin | Asymmetry in extreme local events. |

**Table S1.** Feature name lookup table with original *hctsa* feature names.

| **Feature name** | $\boldsymbol{\rho}$ | $\boldsymbol{p}^{\mathrm{BH}}_{catchaMouse16}$ | $\boldsymbol{p}^{\mathrm{BH}}_{hctsa}$ |
|---|---|---|---|
| `nonlin_autocorr_036` | $-0.69$ | $6.0 \times 10^{-5}$ | 0.014 |
| `increment_ami8` | 0.67 | $1.2 \times 10^{-4}$ | 0.034 |
| `ami3_2bin` | 0.60 | $1.2 \times 10^{-3}$ | 0.41 |
| `outlier_asymmetry` | $-0.60$ | $1.2 \times 10^{-3}$ | 0.43 |
| `walker_crossings` | 0.59 | $1.5 \times 10^{-3}$ | 0.65 |
| `ami3_10bin` | 0.57 | $2.5 \times 10^{-3}$ | 1 |
| `floating_circle` | 0.52 | $9.6 \times 10^{-3}$ | 1 |
| `noise_titration` | 0.35 | 0.30 | 1 |
| `stationarity_min` | 0.29 | 0.63 | 1 |
| `walker_diff` | 0.27 | 0.70 | 1 |
| `floating_circle` | 0.25 | 0.79 | 1 |
| `prediction_scale` | $-0.16$ | 1 | 1 |
| `outlier_corr` | 0.15 | 1 | 1 |
| `dfa_longscale_fit` | 0.078 | 1 | 1 |
| `nonlin_autocorr_112` | 0.062 | 1 | 1 |
| `nonlin_autocorr_035` | $-0.028$ | 1 | 1 |

**Table S2.** Correlations between mouse PV cell density and each of the *catchaMouse16* features.

| **Feature name** | $\boldsymbol{\rho}$ | $\boldsymbol{p}^{\mathrm{BH}}_{catchaMouse16}$ | $\boldsymbol{p}^{\mathrm{BH}}_{hctsa}$ |
|---|---|---|---|
| `dfa_longscale_fit` | 0.52 | 0.014 | 1 |
| `stationarity_min` | $-0.42$ | 0.14 | 1 |
| `walker_diff` | $-0.33$ | 0.62 | 1 |
| `walker_crossings` | $-0.23$ | 1 | 1 |
| `prediction_scale` | 0.21 | 1 | 1 |
| `nonlin_autocorr_035` | 0.21 | 1 | 1 |
| `outlier_asymmetry` | 0.20 | 1 | 1 |
| `floating_circle` | $-0.18$ | 1 | 1 |
| `ami3_2bin` | $-0.17$ | 1 | 1 |
| `nonlin_autocorr_112` | 0.17 | 1 | 1 |
| `ami3_10bin` | $-0.15$ | 1 | 1 |
| `stationarity_floating_circle` | 0.090 | 1 | 1 |
| `noise_titration` | 0.086 | 1 | 1 |
| `increment_ami8` | 0.069 | 1 | 1 |
| `nonlin_autocorr_036` | 0.063 | 1 | 1 |
| `outlier_corr` | $-0.0097$ | 1 | 1 |

**Table S3.** Correlations between mouse VIP cell density and each of the *catchaMouse16* features.

| Feature name | $\rho$ | $p^{\text{BH}}_{catchaMouse16}$ | $p^{\text{BH}}_{hctsa}$ |
|---|---|---|---|
| noise_titration | −0.51 | 0.020 | 1 |
| ami3_10bin | −0.51 | 0.020 | 1 |
| nonlin_autocorr_036 | 0.50 | 0.023 | 1 |
| ami3_2bin | −0.47 | 0.039 | 1 |
| stationarity_min | −0.46 | 0.044 | 1 |
| increment_ami8 | −0.46 | 0.048 | 1 |
| walker_crossings | −0.44 | 0.061 | 1 |
| stationarity_floating_circle | −0.35 | 0.29 | 1 |
| outlier_asymmetry | 0.35 | 0.29 | 1 |
| nonlin_autocorr_112 | −0.28 | 0.60 | 1 |
| nonlin_autocorr_035 | −0.20 | 1 | 1 |
| walker_diff | −0.17 | 1 | 1 |
| outlier_corr | 0.14 | 1 | 1 |
| prediction_scale | −0.073 | 1 | 1 |
| floating_circle | 0.016 | 1 | 1 |
| dfa_longscale_fit | −0.0016 | 1 | 1 |

**Table S4.** Correlations between mouse inhibitory cell density and each of the *catchaMouse16* features.

| Feature name | $\rho$ | $p^{\text{BH}}_{catchaMouse16}$ | $p^{\text{BH}}_{hctsa}$ |
|---|---|---|---|
| nonlin_autocorr_035 | 0.32 | 0.000 21 | 0.083 |
| nonlin_autocorr_112 | 0.32 | 0.000 22 | 0.095 |
| noise_titration | 0.32 | 0.000 22 | 0.16 |
| nonlin_autocorr_036 | 0.32 | 0.000 22 | 0.10 |
| prediction_scale | 0.32 | 0.000 22 | 0.15 |
| ami3_2bin | 0.32 | 0.000 22 | 0.10 |
| ami3_10bin | 0.32 | 0.000 22 | 0.11 |
| floating_circle | −0.31 | 0.000 22 | 0.17 |
| walker_diff | −0.31 | 0.000 23 | 0.27 |
| walker_crossings | −0.30 | 0.000 45 | 0.46 |
| outlier_asymmetry | 0.29 | 0.000 65 | 0.51 |
| stationarity_min | −0.27 | 0.0014 | 1 |
| dfa_longscale_fit | 0.24 | 0.0047 | 1 |
| outlier_corr | −0.24 | 0.0047 | 1 |
| increment_ami8 | 0.23 | 0.0047 | 1 |
| stationarity_floating_circle | 0.17 | 0.025 | 1 |

**Table S5.** Correlations between human *PVALB* expression and each of the *catchaMouse16* features.

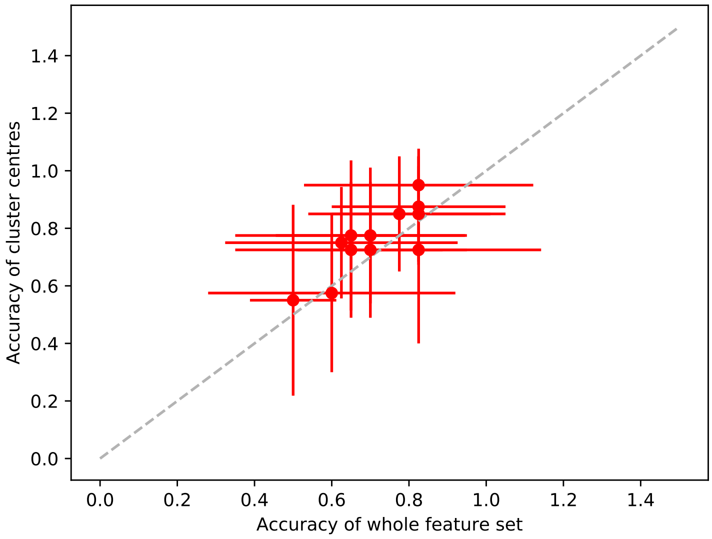


**Figure S1. The centroid features from each feature cluster performs very similar to *catchaMouse16* features.** Mean 10-fold cross validated balanced classification accuracy of the centroids of each cluster (vertical axis) versus *hctsa* (horizontal axis) across the twelve classification tasks. Performance is only marginally improved on the results in Fig. 3. These features are significantly more computationally expensive, but only marginally improve performance. By contrast, *catchaMouse16* proves an efficient yet effective representation of relevant dynamical structure.

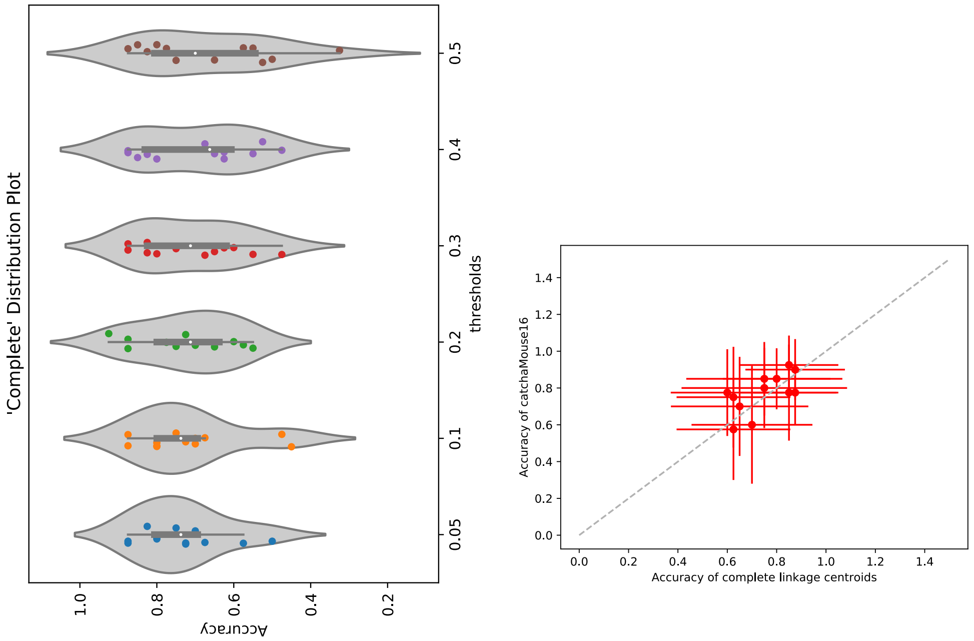


**Figure S2. Comparison of 'average' and 'complete' linkage clustering. Left:** The distribution of balanced accuracies aggregated across all tasks are shown as violin plots for each threshold, $\gamma$ Observe that compared to Fig. 1 the distribution 'complete' performs similarly if not better for $\gamma \geq 0.3$ but worse for $\gamma < 0.3$. **Right:** The accuracy of each task with respect to the Complete cluster centroids (horizontal axis) and the *catchaMouse16* average feature set (vertical axis). A small improvement is observed with the mean accuracy of the complete set 0.75 and the *catchaMouse16* set at 0.77.
